# Extracellular activity of a bacterial protease associated with reduced phage infectivity

**DOI:** 10.1101/2025.09.03.674096

**Authors:** Ehud Herbst, Gael Rosen Blechman, Taya Fedorenko, Sarah Melamed, Gil Amitai, Rotem Sorek

**Affiliations:** Department of Molecular Genetics, Weizmann Institute of Science, Rehovot, Israel

## Abstract

To defend against bacteriophage (phage) infection, bacteria have developed various defense systems, dozens of which were discovered and mechanistically studied recently. To date, almost all defense systems whose mechanisms were deciphered were shown to operate within the bacterial cell. Here we describe a secreted protease from the Actinobacterium *Salinispora mooreana* which, when expressed heterologously in *Streptomyces coelicolor*, reduces titers of two taxonomically related *Siphoviridae* phages. Antiphage effects were maintained when concentrated supernatant from *S. coelicolor* expressing the *Salinispora* protease was added externally to phage-containing medium, even in the absence of bacterial cells, supporting an extracellular mechanism. We further show that phages can escape the antiphage effect of the *Salinispora* protease by mutating a tail-associated protein. The antiphage effect is associated with an increased proportion of phage particles devoid of DNA. Our data suggest antiphage activity of a secreted bacterial protease.

## Introduction

Over the past years, many immune systems that protect bacteria from bacteriophage (phage) infection were discovered and characterized (Doron et al., 2018; Gao et al., 2020; Millman et al., 2022). Various mechanisms of defense were described, such as degradation of phage nucleic acids, regulated death of infected cells, inhibition of protein translation and DNA replication, and depletion of essential metabolites (Georjon and Bernheim, 2023; Tal and Sorek, 2022). Most of the defense systems studied to date operate inside the bacterial cell after the phage has injected its DNA into the cell.

Previous studies have proposed that bacteria might use extracellular proteases to directly inactivate phage particles, but mechanistic studies to directly associate extracellular protease activity to an antiphage effect were limited. Hoque et al showed that incubation of *Vibrio* phages with spent media of *Vibrio cholerae*, exhibiting protease activity, led to decreased phage titers (Hoque et al., 2016). Castillo et al showed that incubation of phages in filtered supernatants from *Vibrio anguillarum* strains led to a reduction of up to 10^3^-fold in phage titers (Castillo et al., 2019). This phage titer reduction was partially reversed by EDTA addition, supporting a potential proteolytic function. However, the identity of the putative proteases responsible for the proteolytic function was not determined (Castillo et al., 2019).

Extracellular proteases have diverse roles in bacterial cell biology. For example, extracellular proteases from *Clostridia* are efficient collagenases; they enable these bacteria to colonize their host and are used clinically for treatment of collagen-related diseases (Eckhard et al., 2011; Hurst et al., 2009; Ramírez-Larrota and Eckhard, 2022). *Streptomyces griseus* trypsin, encoded by the SprT gene, is an enzyme active at the onset of sporulation (Kato et al., 2005). The production of another extracellular protease, with chymotrypsin-like activity in *Streptomyces exfoliates*, was associated with mycelium formation and suggested to be important for the acquisition of proteinaceous nitrogen sources (Kim and Lee, 1995).

In this study, we report the identification of a bacterial gene coding for a putative extracellular protease from the Actinobaterium *Salinispora mooreana*, which exhibits an antiphage activity. *Streptomyces coelicolor* that heterologously expresses this protease can be infected by *Streptomyces* phages, but following replication and prolonged incubation, some phages lose infective titers by up to three orders of magnitude. The effect is specific for two homologous phages among a collection of *Streptomyces coelicolor* phages tested. The antiphage effect of the *Salinispora* protease is maintained when supernatant of *S. coelicolor* expressing the *Salinispora* protease is added externally to *S. coelicolor* cells not expressing the *Salinispora* protease, or incubated with phages in the absence of bacterial cells, supporting an extracellular activity of the secreted protease. By isolating mutant phages that can escape the antiphage effect of the *Salinispora* protease we identify a phage gene encoding for a structural protein that is repeatedly mutated in escaper phages. Our electron microscopy data suggest that the antiphage effect of the *Salinispora* protease is associated with counterproductive DNA ejection from affected phages.

## Results

While examining putative gene cassettes predicted to encode antiphage pathways in bacteria from the genus *Salinispora*, we noticed a gene coding for a protease that was sometimes present next to various defense systems in these bacteria (Figure 1A). This gene caught our attention because its N-terminus was predicted to harbor a signal peptide, suggesting that it encodes for a secreted protease with extracellular proteolytic activity (Figure 1B). We decided to test the protease-encoding gene from *Salinispora mooreana* for antiphage defense. We chose *Streptomyces coelicolor* as the heterologous host to express this gene, because of its frequent use as a heterologous host for expressing genes derived from species of the *Actinomycetota* phylum, and because a variety of phages that infect this species were described (Gomez-Escribano and Bibb, 2011; Hardy et al., 2020; Russell et al., 2017; Zhang et al., 2019a).

**Figure 1.**
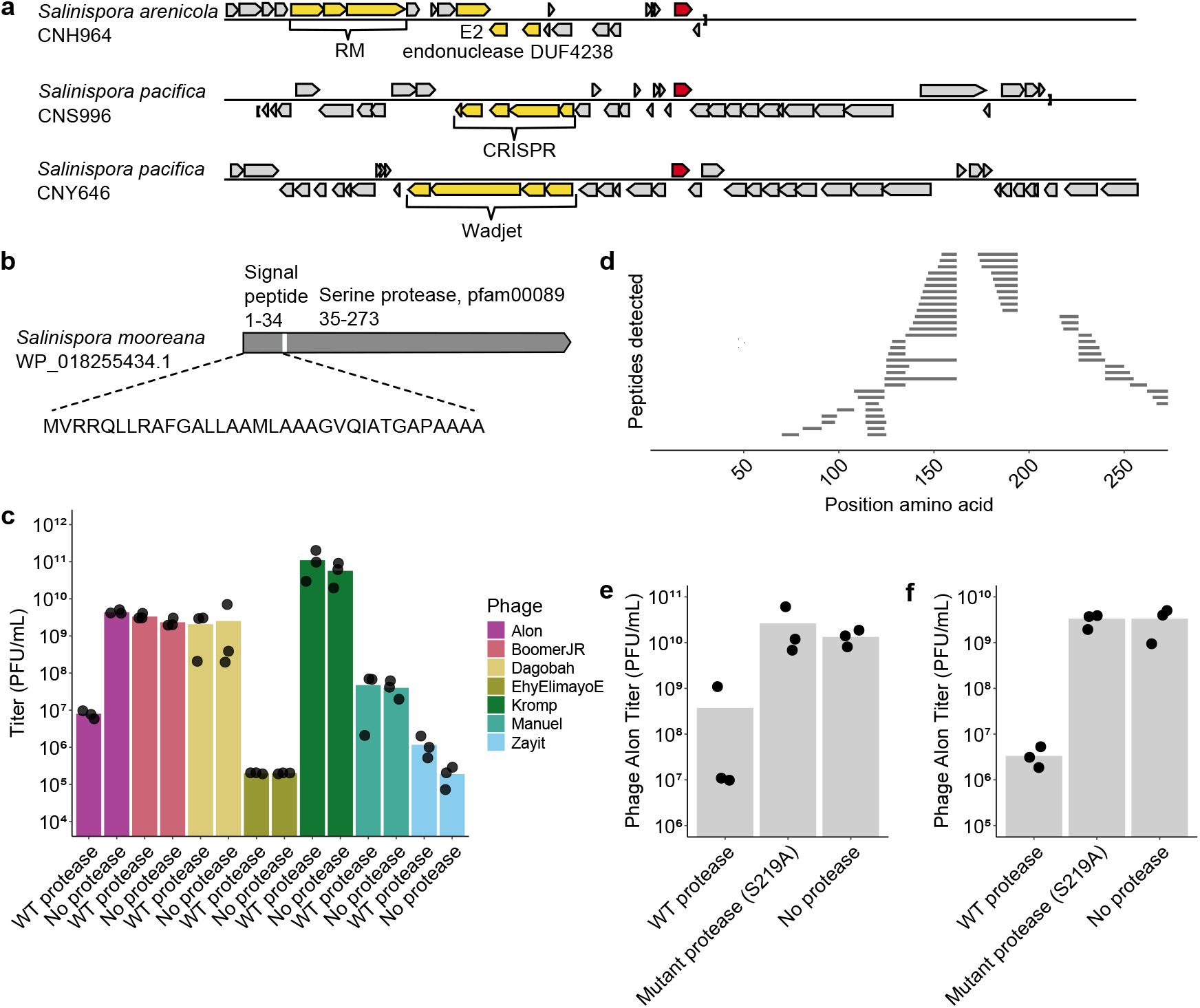
A secreted protease associated with reduced phage titers. **a**. Genomic neighborhoods of a group of predicted secreted proteases (red) in *Salinispora* species. Genes known to be involved in bacterial antiphage defense are in yellow; other genes in grey. **b**. Position of a signal peptide in the *Salinispora* protease as predicted by the SignalP-6.0 software (Teufel et al., 2022). **c**. Titers of seven phages following infection with *S. coelicolor* expressing the WT *Salinispora* protease or an apramycin resistance gene used as a negative control. Phages were initially added at multiplicity of infection (MOI) of 0.05, and harvested from the culture two days from the onset of infection. Phages were then counted by plating on an indicator strain. Bars show the averages of three biological replicates with individual data point overlaid. **d**. Mass spectrometry (MS) analysis indicates presence of the protease in supernatants. Shown are non-redundant peptides (n=60) detected using MS. *Streptomyces coelicolor* expressing the WT *Salinispora* protease was grown for 46 h, and the supernatant was harvested and filtered. Samples were digested with trypsin and were then subjected to protein MS. **e**. Titers of phage Alon following infection in the presence of protease-containing supernatant. WT or mutant protease-expressing *S. coelicolor* cells or cells not expressing the protease were grown for two days in the absence of phage, and supernatant was collected, filtered and concentrated. This supernatant was added to an infection experiment in liquid culture with *S. coelicolor* cells not expressing the protease and phage Alon at MOI=0.1. Phages were then harvested from the culture two days after infection. Phage titer counts shown are averages of three biological replicates with individual data point overlaid. **f**. Titers of phage Alon following infection with *S. coelicolor* expressing the WT *Salinispora* protease (WT), a catalytically dead protease mutant or an apramycin resistance gene instead of the protease as control. Phages were initially added at MOI=0.1, and harvested from the culture two days from the onset of infection. Bars show the averages of three biological replicates with individual data point overlaid.

To test whether the *Salinispora* protease has an antiphage effect, we performed infection experiments in liquid culture. *S. coelicolor* expressing the *Salinispora* protease, as well as control *S. coelicolor* not expressing the protease, were subjected to infection by a set of seven diverse *S. coelicolor* phages, some of which obtained from culture collections while others were isolated by us. We followed the dynamics of phage replication by taking samples from the liquid culture two days following phage addition, and used plaque assays on an indicator protease-less *S. coelicolor* to count phages. Following prolonged incubation for two days, the infective titers of phage Alon, but not any of the other phages, was reduced by three orders of magnitude when infecting the protease-expressing strain (Figure 1C).

The *Salinispora mooreana* protease was predicted by the SignalP-6.0 software to encode an N-terminal twin arginine translocation signal peptide of 34 amino acids with 99.9% likelihood (Teufel et al., 2022). We verified by protein mass spectrometry (MS) that the *Salinispora* protease is indeed found in the supernatant from a culture of *S. coelicolor* expressing the *Salinispora* protease (Figure 1D, Table S1). Peptides spanning the N-terminal signal peptide were depleted in these data, further supporting the hypothesis that this protease is secreted and the signal peptide is cleaved in the process (Figure 1D).

To test the hypothesis that the *Salinispora mooreana* protease exerts an antiphage activity extracellularly, we added concentrated supernatants from *S. coelicolor* expressing the *Salinispora* protease to media in which we performed infection experiments with phage Alon and *S. coelicolor* not expressing the protease. The results of this experiment demonstrated supernatant-dependent reduction in titers of phage Alon (Figure 1E).

To further substantiate that the proteolytic activity of the *Salinispora* protease is responsible for phage titer reduction, we substituted serine residue 219 in the protease, predicted to participate in the catalytic triad of the active site (Matthews et al., 1967; Morrison, 2021; Yousef et al., 2004), to alanine. Indeed, our data show that phage Alon infecting *S. coelicolor* cells that express the catalytically dead S219A variant does not exhibit titer reduction (Figure 1F). In further support of this result, adding concentrated supernatant from *S. coelicolor* expressing the catalytically dead S219A protease to media where infection experiments where performed with phage Alon and *S. coelicolor* lacking the protease, did not lead to phage titer reduction (Figure 1E).

To gain further insight into the mechanism underlying the protease effect on phage titers, we attempted to isolate phage mutants that escape the effect of the protease. Such escaper mutants were previously shown useful for predicting the specificity determinants of bacterial defense systems (Stokar-Avihail et al., 2023). We collected phages from an infection experiment in liquid culture of *S. coelicolor* expressing the *Salinispora* protease and subjected them to two additional rounds of infection with bacteria expressing the *Salinispora* protease. Isolated escapers showed a partial or complete resistance to the antiphage effect of the *Salinispora* protease (Figure 2A). We have not observed such phenotype in single plaques isolated from phage survivors of infections in the presence of mutant *Salinispora* protease (Figure 2A).

**Figure 2.**
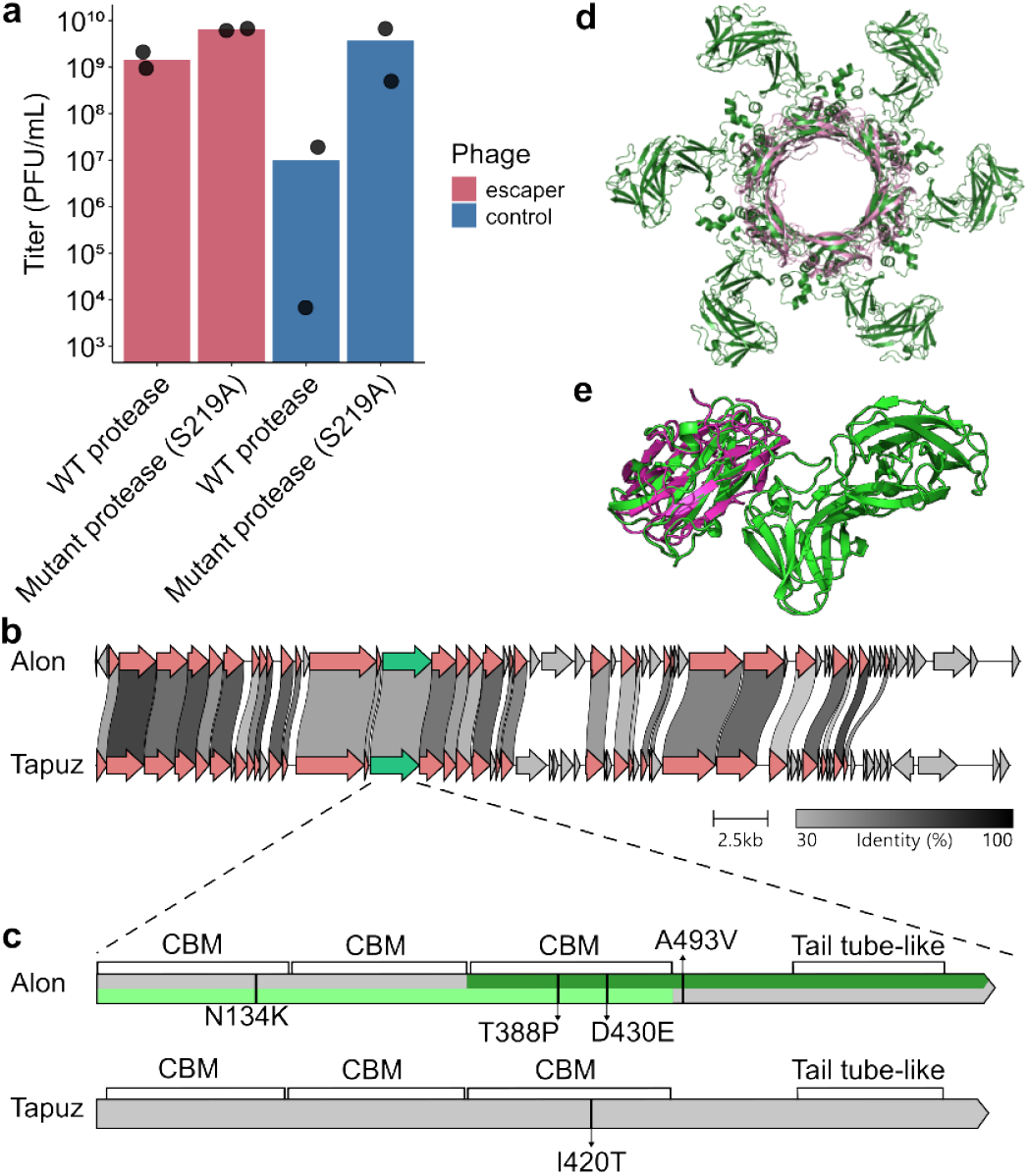
Phages escape the effect of the protease via mutations in a tail-associated gene. **a**. Phage Alon variants selected on bacteria expressing the WT *Salinispora* protease (red) or catalytically dead protease (blue). Results shown for a representative escaper mutant and control are averages of two biological replicates (two separately isolated escaper phages with the same genotype) with individual datapoints overlaid. Data for all phage Alon mutants that were isolated and sequenced are presented in Figure S2 and Table S2. **b**. Genome comparison of phage Alon and phage Tapuz. Amino acid sequence similarity in the range of 30%-100% is marked by grey shading. The gene mutated in all escapers is marked in green. Other genes with over 30% identity between the two phages are marked in red. Visualized using clinker (Gilchrist et al., 2021). **c**. Position of point mutations on the phage tail-associated protein, identified in phage mutants that escaped the effect of the protease. Domains are annotated according to a FoldSeek homolog search from the AlphaFold2 predicted structure (Jumper et al., 2021; van Kempen et al., 2023), with CBM indicating carbohydrate binding module. Dark and light green mark the portion of the protein modeled in panels 2D and 2E, respectively. **d**. The C terminus of the protein mutated in phage Alon escapers is predicted by Alphafold to form hexamers (model ipTM=0.79). The hexameric structure (dark green, amino acids 317-759) is structurally homologous (amino acids 589 through 716) to the tail tube protein of phage T4 (PDB code 5w5f, pink, one hexamer out of three hexamers presented in 5w5f is plotted for clarity). **e**. AlphaFold2 prediction of the N-terminus of the tail-associated protein (light green, amino acids 1 through 486) aligned with a homologous carbohydrate binding module (PDB code 3k4z, purple, amino acids 1 through 176).

We sequenced the genome of 5 isolated escaper phages that were partially or wholly resistant to the antiphage effect of the *Salinispora* protease. As a control we sequenced the genomes of 3 phages selected in an infection experiment in liquid culture of bacteria expressing mutant protease. All 5 escapers isolated on strains expressing the WT protease had a mutation in the same phage gene (Figure 2B, Figure 2C), but none of the control phages had mutations in that gene (Table S2). Four of the escaper phages had an additional deletion that was also observed in control phages selected on bacteria expressing a mutant protease, suggesting that this deletion is unrelated to the escape phenotype (Table S2).

To test the robustness of the immune evasion phenotype of the escaper phages, we repeated this process also for a related phage called Tapuz, which shows 70% homology to phage Alon over 42% of its nucleotide sequence (Sayers et al., 2025), and which we also found sensitive to the effect of the *Salinispora* protease (Figure S1). Two Tapuz phage mutants that can escape protease-mediated defense were isolated, and in both cases these phages exhibited a mutation in an orthologous gene in phage Tapuz, which shares 46% sequence identity with the protein mutated in the phage Alon escapers (Figure 2B, 2C). One of the escaper phages (escaper no. 6) did not contain any other mutation except for the mutation in the indicated gene, showing that this mutation is sufficient for the escaper phenotype (Table S2).

We predicted the structure of the mutated phage protein using AlphaFold2 in monomer and multimer forms (Jumper et al., 2021) and ran a structural homology search using Foldseek (van Kempen et al., 2023). The C-terminus of the protein (amino acids 589 through 716 in phage Alon) showed structural homology to the tail tube protein of phage T4 (Zheng et al., 2017), suggesting that this protein may function as a tail-associated protein in phage Alon. Similar to the T4 tail tube, the AlphaFold2 model for the C-terminus of the phage Alon tail tube predicted high confidence (ipTM=0.79) hexameric structure of a similar diameter to the phage T4 hexameric protein (Figure 2D). The N-terminus of the predicted tail tube protein of phage Alon exhibits three consecutive domains that are structurally similar to each other as well as to carbohydrate binding modules (Figure 2E).

Our data show that phages mutated in a predicted tail-associated protein can escape the effect of the *Salinispora* protease. To test the hypothesis that the antiphage effect of the *Salinispora* protease results from direct interaction with the phage particle, we incubated purified phage particles with concentrated supernatant from bacteria expressing the *Salinispora* protease. Phages incubated with the supernatant alone for one day showed 10-fold less infectivity as compared to phages incubated with a supernatant from bacteria expressing the mutated protease (Figure 3A). As with the antiphage effect in infection experiments in liquid culture, this bacteria-independent antiphage effect of the *Salinispora* protease was not observed in phages not homologous to phage Alon (Figure 3A).

**Figure 3.**
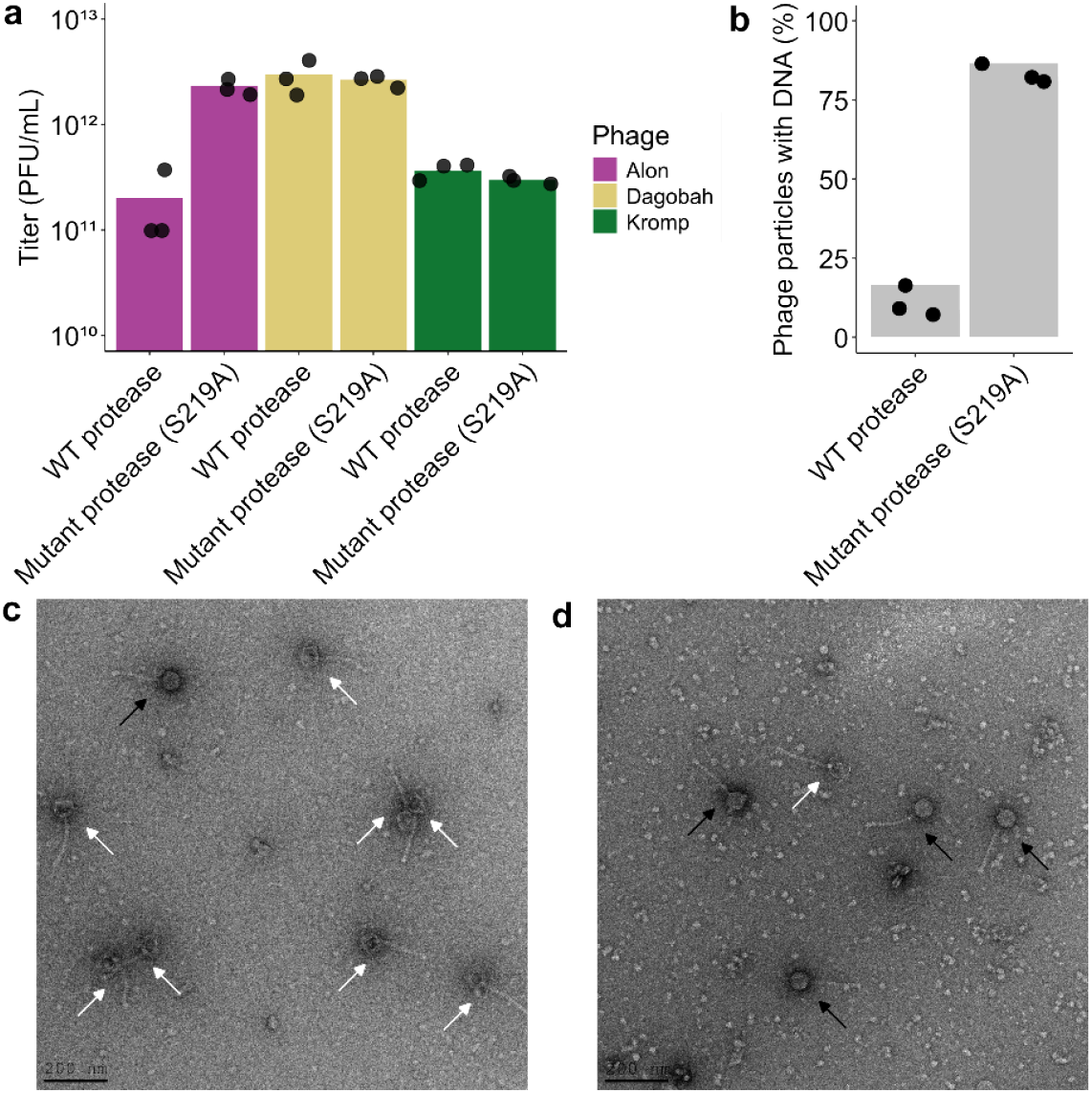
The *Salinispora* protease is associated with Phage Alon titer reduction and DNA loss. **a**. Incubation of phage particles with protease-containing supernatant results in titer loss. Phages Alon, Dagobah and Kromp were purified using CsCl gradient and incubated with concentrated supernatant from *S. coelicolor* cells expressing WT or catalytically dead *Salinispora* protease. Phage titers shown are averages of three biological replicates with individual data points overlaid. **b**. *Salinispora* protease-dependent DNA-loss in phage Alon when incubated with concentrated supernatant from *S. coelicolor* cells expressing WT or catalytically dead *Salinispora* protease. Particles were imaged under transmission electron microscopy and DNA content was recorded based on the observed density of the phage head. Graphs represent average percent of phages particles with DNA to total phage particles in counts of 300 particles per replicate. Shown is average of three replicates with individual data points overlaid. **c**. *Salinispora* protease-dependent DNA loss in phage Alon incubated with concentrated supernatant from *S. coelicolor* cells expressing WT *Salinispora* protease. Representative electron microscopy images of phages following treatment with concentrated supernatant from *S. coelicolor* expressing WT (**c**, left) or catalytically dead S219A *Salinispora* protease (**d**, right). Black arrows point to phage particles with DNA and white arrows point to DNA-less “ghost” phage particles. Additional electron microscopy fields are presented in Figure S3.

We next imaged the particles of phage Alon by transmission electron microscopy. Following incubation of the phages with concentrated supernatant obtained from protease-expressing bacteria, we observed that 90% of the phages contained “empty” heads, suggesting that these phages lost their DNA content (Figure 3B-D). In contrast, Alon phages incubated with supernatant obtained from bacteria expressing the mutated protease mostly exhibited DNA-containing heads (Figure 3B-D, Figure S3). These data suggest that phages incubated with protease-containing supernatant tend to have their DNA ejected, explaining the reduction in infectivity for these phages.

## Discussion

In this study we demonstrate that a protease predicted to be secreted can affect the outcome of phage infection. When expressed in *S. coelicolor*, this *Salinispora*-originating protease is able to reduce infective titers of phages Alon and Tapuz after prolonged incubation, and we show that this activity is localized outside of the bacterial cell. The loss of phage infective titers was associated with increased ratios of phages that lost their DNA, suggesting that the protease causes loss of the DNA from affected phages. We furthermore found that phages mutated in a likely tail-associated protein can escape the effect of the protease.

One can envision several hypothetical mechanistic scenarios that can explain our observations. Under one hypothesis, cleavage of the structural protein in the phage tail leads to premature DNA ejection independent of the presence of bacteria, resulting in the observed DNA-less particles in the electron microscope. Within this hypothesis, escaper phages are resistant either to the direct proteolytic effect of the protease on the tail-associated protein or are able to maintain their DNA unejected even when their tail-associated protein is subjected to proteolysis.

An alternative hypothesis involving proteolysis is that the *Salinispora* protease cleaves off an extracellular portion of the bacterial protein used by the phage as a receptor. Within this hypothesis, the cleaved portion of the receptor protein is present in the supernatant derived from cells expressing the *Salinispora* protease, and the phage might bind it, thereby stimulating DNA ejection. Escaper phage mutants may have a reduced affinity to the phage receptor and require increased receptor concentration to lead to binding and DNA ejection. Phage T5, for example, is known to release its DNA *in vitro* in the presence of its bacterial receptor FhuA (Leforestier and Livolant, 2010).

Our study has several limitations. First, it is possible that the primary role of the protease does not involve phage defense when expressed in its original *Salinispora* host. However, studies in the original bacteria encoding this protease are limited due to the slow growth rate of this marine bacterium in laboratory settings (Román-Ponce et al., 2020), as well as the absence of genetic systems and the lack of known phages for this host. While the protease we studied is located in variable genomic environments in *Salinispora* species, homologs of this protease in other bacteria do not tend to be enriched next to known defense systems in bacterial genomes, implying that this family of proteases have roles other than phage defense. It is therefore possible that the effect we observed when expressing this protease in *S. coelicolor* is a side effect of the real biological activity of the protease.

Another limitation of this study is that we worked with supernatants from protease-expressing cells rather than with a purified protease. While we demonstrated that supernatants contain the secreted protease, and that the antiphage activity we found in the supernatants is dependent on an intact protease active site, we cannot rule out that the activity we observed involves other factors. Unfortunately, we have not managed to purify this protease from *E. coli* cells. Future studies with purified proteases can examine whether the protease alone has a direct effect on premature DNA ejection from phage particles.

A puzzling observation in our study is that phage Alon can readily infect and replicate on *S. coelicolor* cells even when these cells express the *Salinispora* protease (Figure S4). The reduction in infective titers is only observed after the phages, which already replicated, are further incubated with the cells (or the supernatant derived from these cells) for a prolonged period of two days. This can be explained in several ways. First, it is possible that the protease activity is very slow, and only affects the phages after long incubation. Second, it is possible that the antiphage activity necessitates another factor that is produced by *Streptomyces* cells only in a late developmental stage. It is known that *Streptomyces* species produce some secondary metabolites only after 2-3 days from spore germination (Manteca et al., 2008), and in some cases even longer periods are required for the production of secondary metabolites (Barbuto Ferraiuolo et al., 2021; Manteca et al., 2008).

Most bacterial defense systems investigated to date were heterologously expressed in *E. coli* or *B. subtilis* and originated from members of their respective phyla (Doron et al., 2018; Gao et al., 2020; Millman et al., 2022). Recent studies showed that using expression hosts from the *Actinomycetota* phylum, and specifically *Streptomyces* bacteria, can lead to the discovery of new concepts in antiphage defense (Kronheim et al., 2018; Luthe et al., 2023; Mordret et al., 2025; Shomar et al., 2024). Our study is an additional demonstration of the benefit of using *Streptomyces* to study bacterial defenses. As *Actinomycetota* often live in complex ecological environments and exhibit multicellularity characteristics, it might make sense that these bacteria would involve extracellular antiphage activity as part of their defensive arsenals. Further studies with *Streptomyces* may find additional such extracellular mechanisms in the future.

## Supplementary information

**Figure S1.**
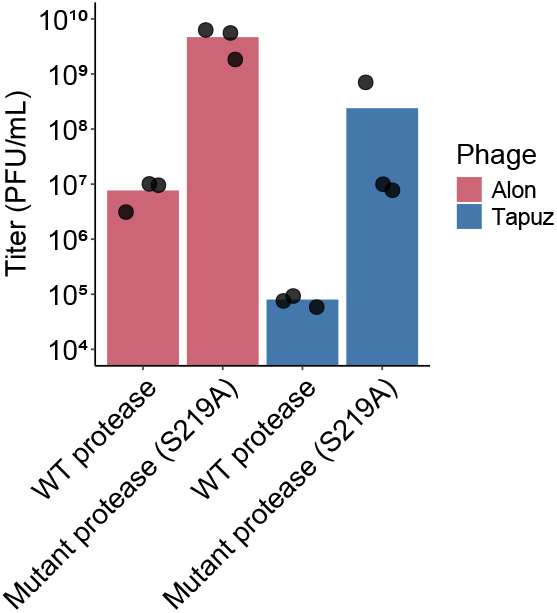
Titers of phages Alon and Tapuz following infection with *S. coelicolor* expressing the WT *Salinispora* protease (WT) or a catalytically dead mutant of the protease. Phages were initially added at MOI=0.05, and harvested from the culture two days from the onset of infection. Bars show the averages of three biological replicates with individual data point overlaid.

**Figure S2.**
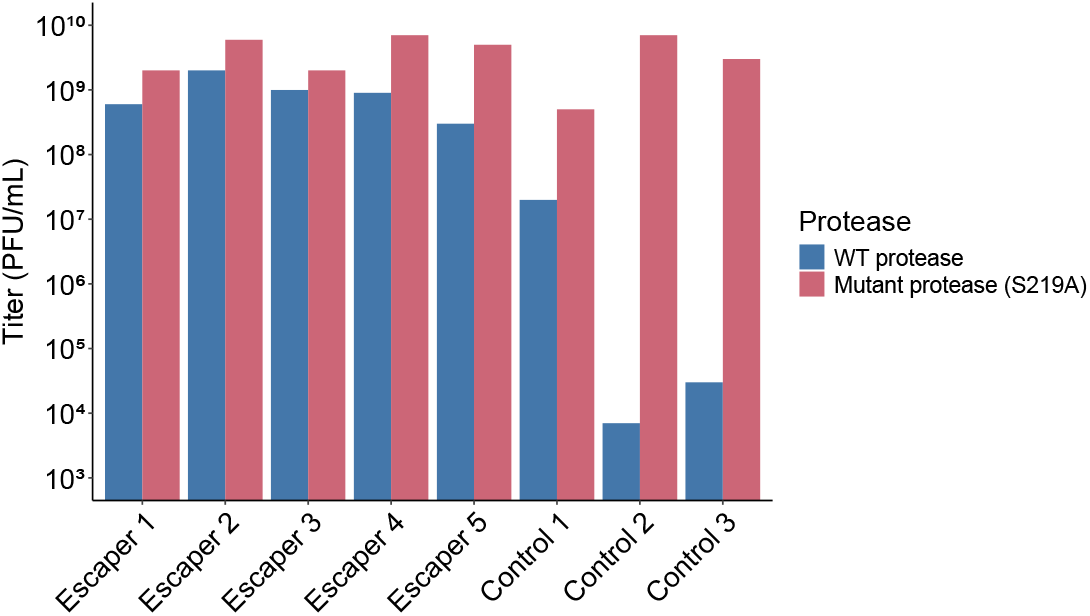
Phage Alon variants selected on bacteria expressing the WT *Salinispora* protease (Escaper) or catalytically dead protease (Control). Shown data are results obtained for five escaper phages and three control phages. Genotypes for all mutants that were isolated and sequenced are presented in Table S2. Data for Escapers 2 and 4, and for Control phages 1 and 2, are also presented in Figure 2A.

**Figure S3.**
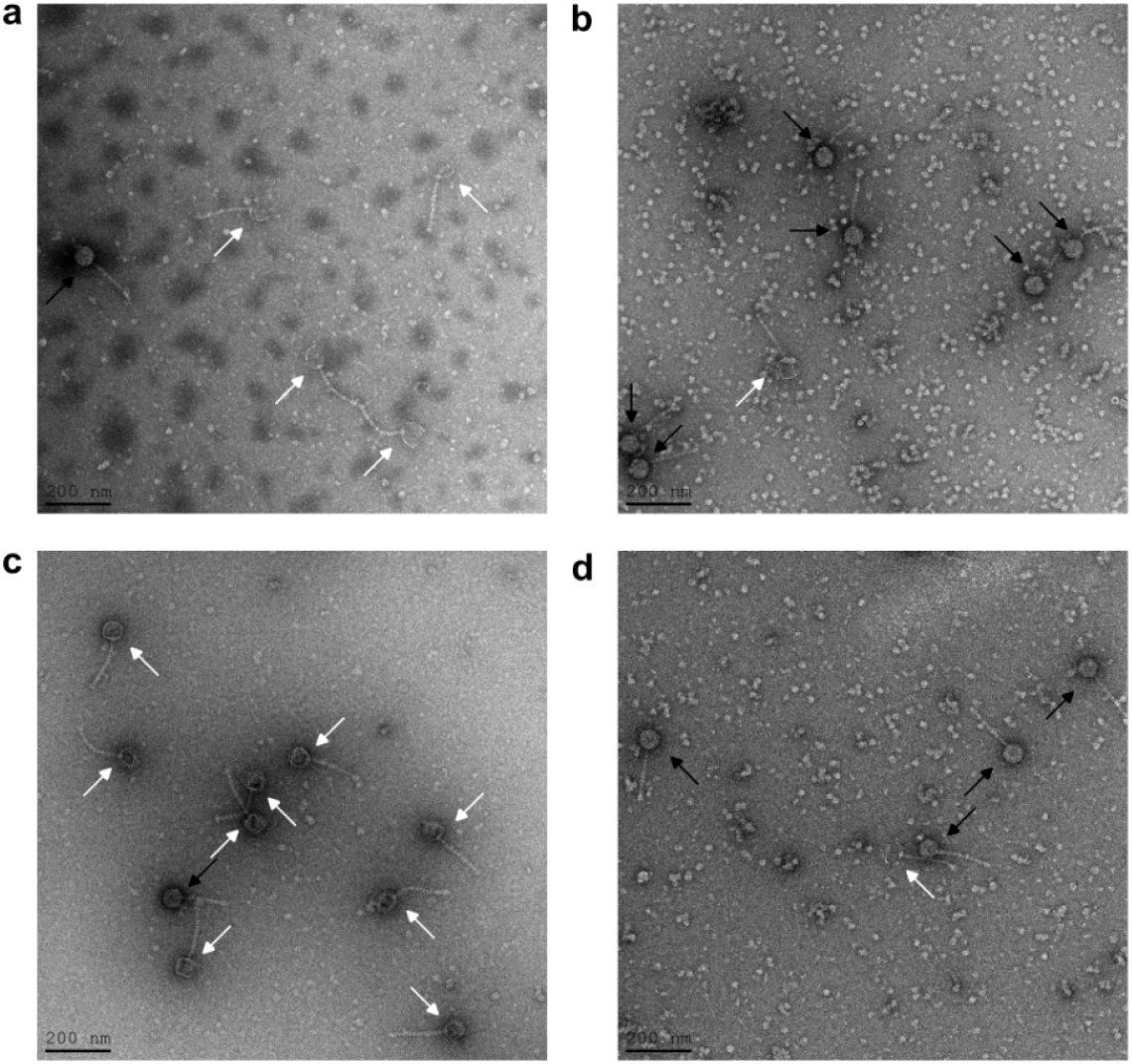
*Salinispora* protease-dependent DNA loss in phage Alon incubated with concentrated supernatant from *S. coelicolor* cells expressing WT *Salinispora* protease but not mutant protease. Additional representative electron microscopy images of phages following treatment with concentrated supernatant from *S. coelicolor* expressing WT (**a, c**, left) or catalytically dead S219A *Salinispora* protease (**b, d**, right). Black arrows point to phage particles with DNA and white arrows point to DNA-less phage “ghost” particles.

**Figure S4.**
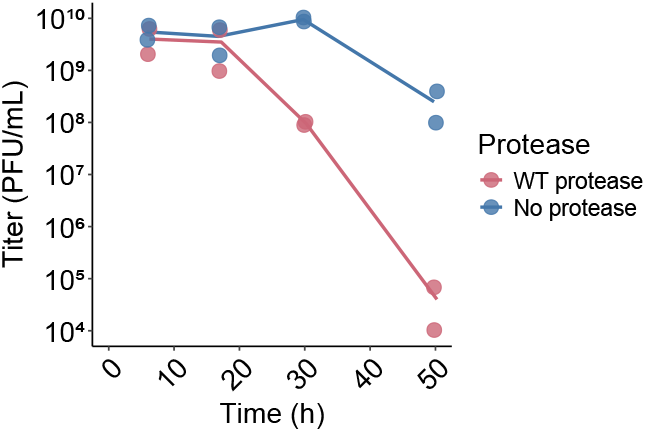
Time course experiment with phage Tapuz showing how phage titers change over time in an infection experiment of *S. coelicolor* encoding the WT *Salinispora* protease (red) or a negative control not encoding the protease (blue). One of the two strains encoding the protease (red), encodes it within a 1.4 kb genomic fragment derived from *Salinispora mooreana* (NCBI Reference sequence: NZ_KB900614.1; range: 4542153 to 4543551), while the other encodes it within a larger genomic fragment of 15.2 kb taken from the same strain (NCBI Reference sequence: NZ_KB900614.1; range: 4542146 to 4557358). One of the control strains (those not encoding the protease, blue), encodes an apramycin resistance gene instead of the genomic fragment containing the protease (pCAP03-acc(3)IV, Addgene 69862; Range: 700 to 2075), while the other encodes genomic fragments from *Streptomyces anulatus* instead of the genomic fragment containing the protease (GenBank: HM038106.1; ranges: 576 to 625 and 47697 to 47746, separated from each other by a PmeI restriction site). Phages were initially added at a concentration of 3×10^5^ PFU/mL, MOI=0.01 and harvested from the culture after 6 h, 17 h, 30 h and 50 h from the onset of infection. Lines show the averages of two strains encoding or not encoding the protease with individual data point overlaid.

**Table S2.** Phages used in this study.

| Phage number | Escaper number | Phage name | Host bacteria amplified on | Mutations (escaper phages) | Comments | Source |
| --- | --- | --- | --- | --- | --- | --- |
| 1 |  | Tapuz | M145 |  |  | This study |
| 2 |  | Zayit | M145 |  |  | This study |
| 3 |  | Alon | M145 |  |  | This study |
| 4 |  | Boomer | M145 |  |  | Phages DB |
| 5 |  | Manuel | *S. lividans* 1326 |  |  | Phages DB |
| 6 |  | EhyElimayoE | *S. lividans* 1326 |  |  | Phages DB |
| 7 |  | Dagobah | M145 |  |  | DSMZ |
| 8 |  | Kromp | M145 |  |  | Phages DB |
| 9 | 1 | Alon D430E | 5) WT protease | C→A in position 14560 and Δ115 bp in 33665 | Escaper mutant in tail-associated protein | This study |
| 10 | 2 | Alon T388P | 5) WT protease | A→C in position 14432 and Δ115 bp in 33665 | Escaper mutant tail-associated protein | This study |
| 11 | 3 | Alon N134K | 5) WT protease | C→A in position 13672 and Δ115 bp in 33665 | Escaper mutant in tail-associated protein | This study |
| 12 | 4 | Alon T388P | 5) WT protease | A→C in position 14432 and Δ115 bp in 33665 | Escaper mutant in tail-associated protein | This study |
| 13 | 5 | Alon A493V | 5) WT protease | C→T in position 14748 and C→G in 33731 | Escaper mutant in tail-associated protein | This study |
| 14 |  | Alon control escaper | 6) Mutant protease | Δ115 bp in position 33665 | Control escaper isolated on mutant S219A protease | This study |
| 15 |  | Alon control escaper | 6) mutant protease | Δ115 bp in position 33665 | Control escaper isolated on mutant S219A protease | This study |
| 16 |  | Alon control escaper | 6) Mutant protease | Δ115 bp in position 33665 | Control escaper isolated on mutant S219A protease | This study |
| 17 | 6 | Tapuz I420T | 5) WT protease | T→C in position 13964 | Escaper mutant in tail-associated protein | This study |
| 18 | 7 | Tapuz I420T | 5) WT protease | T→C in position 13964 and T→C in 28891 | Escaper mutant in tail-associated protein | This study |
| 19 |  | Tapuz control escaper | 6) Mutant protease | none | Control escaper isolated on mutant S219A protease | This study |
| 20 |  | Tapuz control escaper | 6) Mutant protease | none | Control escaper isolated on mutant S219A protease | This study |

**Table S3.** Bacterial strains used in this study.

| Strain number | Strain description | Parent strain | Integrative plasmid | Source |
| --- | --- | --- | --- | --- |
| 1 | *S. coelicolor* M145 |  | - | Mervyn J. Bibb, John Innes Centre |
| 2 | *S. coelicolor* M1146 |  | - | Mervyn J. Bibb, John Innes Centre |
| 3 | *S. lividans* 1326 |  | - | DSMZ 46482 |
| 4 | *E. coli* ET12567 |  | - | ATCC BAA-525 |
| 5 | WT *Salinispora* protease in M1146 | M1146 | 15_2517459620_Gib_in_pCAP03 | This study |
| 6 | Mutant *Salinispora* protease S219A in M1146 | M1146 | 15_2517459620_S219A_Gib_in_pCAP03 | This study |
| 7 | pCAP03 (addgene) in M1146 | M1146 | pCAP03 | This study |
| 8 | WT *Salinispora* protease in M1146 within a larger *Salinispora* genome context (15.2 kb) | M1146 | 15_in_pCAP03 | This study |
| 9 | 0.1 kb genome fragment from *Streptomyces anulatus* in M1146 | M1146 | 3_actinomycin-50-bp-homo-capture-vector-pCAP03 | This study |

**Table S4.** Plasmids used in this study.

| Plasmid number | Plasmid name | Plasmid description | Host strain | Source |
| --- | --- | --- | --- | --- |
| 1 | pRK2013 |  | ET12567 | DSMZ 5599 |
| 2 | pCAP03 |  | ET12567 | Bradley Moore, UCSD |
| 3 | 15_2517459620_Gib_in_pCAP03 | WT *Salinispora* protease in integrative plasmid pCAP03 | ET12567 | This study |
| 4 | 15_2517459620_S219A_Gib_in_pCAP03 | mutant *Salinispora* protease S219A in integrative plasmid pCAP03 | ET12567 | This study |
| 5 | 15_in_pCAP03 | WT *Salinispora* protease within a wider *Salinispora mooreana* genomic context (15.2 kb) in integrative plasmid pCAP03 | ET12567 | This study |
| 6 | 3_actinomycin-50-bp-homo-capture-vector-pCAP03 | Acceptor vector for the actinomycin pathway containing 50 bp homology to the edges of the pathway in *Streptomyces anulatus* in integrative plasmid pCAP03 | ET12567 | This study |

## Materials and Methods

Spore stocks of *Streptomyces coelicolor* and *Streptomyces lividans* strains were prepared as previously described (Shepherd et al., 2010). Conjugation and genome integration of integrative plasmids into *Streptomyces coelicolor* was performed as previously described (Zhang et al., 2019b), with the following modification: pRK2013 (Figurski and Helinski, 1979) was used as the helper conjugation plasmid instead of pUB307. Phage infection experiments in liquid culture were performed in DNB MMC media, (Difco nutrient broth, supplemented with 10 mM MgCl_2_, 0.1 mM MnCl_2_ and 8 mM CaCl_2_) at 30 °C in 15 mL culture tubes shaking at 350 rpm, unless specified otherwise. Phage sequence homology search was performed using blastn with the default search parameters (Sayers et al., 2025). Protein structure prediction from sequence was performed using AlphaFold2 (Jumper et al., 2021). Protein structural homolog search was performed using Foldseek (van Kempen et al., 2023). Protein structures were visualized using PyMOL (Schrodinger, 2015).

### Phage isolation from soil

Phages Alon, Tapuz and Zayit were isolated from soil following previously reported protocols (Kronheim et al., 2018), with modifications as described hereby: In a 15 mL culture tube, *S. coelicolor* M145 was added at a concentration of 5×10^6^ CFU/mL to 5 mL of DNB with 0.5% glucose and 4 mM CaCl_2_. 2 mL of soil were added as well and samples were incubated at 30 °C overnight. Supernatant was collected by two rounds of centrifugation followed by filtration. The supernatant was serially diluted and plated on DNB to obtain single plaques as described below: 100 μL spore suspension of *S. coelicolor* M145 (10^8^ CFU / mL) were added to 15 mL screw cap tubes. 5 mL of DNB MMC with 0.5% agar at 50 °C were then added and the resulting phage/spore suspension was mixed and plated on 100 × 15 mm plates with 20 mL DNB 2% agar. The plates were incubated overnight at 30 °C. Single plaques were picked with a pipette tip into 120 μL phage buffer. Two more rounds of plating for single plaque and picking ensued and then phages were amplified as described below, under ‘phage stocks preparation from single plaques’, to obtain the phage stock. Phage DNA sequencing was performed as previously described (Stokar-Avihail et al., 2023).

### Phage stocks preparation from single plaques

To prepare phage stocks from single plaques, phage stocks were serially diluted, 20 μL into 180 μL, and added to a 100 μL spore suspension of *S. coelicolor* M145 (10^8^ CFU / mL) in 15 mL screw cap tubes. 5 mL of DNB MMC with 0.5% agar at 50 °C were then added and the resulting phage/spore suspension was mixed and plated on 100 × 15 mm plates with 20 mL DNB 2% agar. The plates were incubated overnight at 30 °C followed by incubation at 25 °C until single plaques were visible, if necessary. Single plaques were picked with a pipette tip into 120 μL phage buffer.

To obtain confluent lysis, single plaque suspensions in phage buffer were serially diluted, 20 μL into 180 μL, in phage buffer and added to 100 μL spore suspension of *S. coelicolor* M145 (10^8^ CFU / mL) in 15 mL screw cap tubes. 5 mL of DNB MMC 0.5% agar at 50 °C were then added and the resulting phage/spore suspension was mixed and plated on 100 × 15 mm plates with 20 mL DNB 2% agar.

Phage stocks were prepared from confluent lysis plates by scraping the 5 mL 0.5% agar layer onto 6 mL DNB in 15 mL screw cap tubes, vortexing and shaking intermittently for 1 h at room temperature, followed by 15 min centrifugation at 3200 *g* at 4 °C. The supernatant was then filtered through 0.2 μ filters and phage concentration was measured by plaque assay, as described below, under ‘phage titer measurement by plaque assay’.

### Infection experiment of *Streptomyces coelicolor* with phages in liquid culture

Spores of *Streptomyces coelicolor* encoding WT *Salinispora* protease, mutant S219A *Salinispora* protease or no *Salinispora* protease (Table S3, strains 5-7) were added in biological triplicates to 750 μL of DNB MMC in culture tubes of 15 mL at a concentration of 3×10^7^ CFU/mL, along with relevant phage in multiplicity of infection (MOI) of 0.1 and incubated for 45 h, unless otherwise indicated. Following incubation, phages were harvested by pelleting bacterial cells at 3200 g at 4 °C for 30 min, followed by filtration through 0.2 μ cellulose acetate filters. Phage titer was then measured by plaque assay as described blow.

### Phage titer measurement by plaque assay

Phage stocks were serially 10× diluted, 20 μL to 180 μL, in phage buffer (50 mM Tris pH 7.4, 100 mM MgCl_2_, 10 mM NaCl), to obtain 10^0^-10^−7^ dilutions. A square petri dish of *Streptomyces coelicolor* M145 was prepared by mixing 30 mL of 50 °C DNB MMC medium with 0.5% agar with 10^6^ CFU of M145 spores and left to cool for 1 h. Onto that plate, 10 μL of each phage dilution were dropped, left to dry for 1 h and placed in 30 °C overnight, followed by incubation at 25 °C until single plaques were visible, if necessary. Phage titer was calculated by counting individual plaques in the highest concentration dilution where single plaques are countable and multiplying by the corresponding dilution factor.

### Supernatant harvest from *Streptomyces coelicolor* cultures

Spores of *Streptomyces coelicolor* encoding WT *Salinispora* protease, mutant S219A *Salinispora* protease or no *Salinispora* protease (Table S3, strains 5-7) were added in biological triplicates to 6 mL of DNB MMC at a concentration of 3×10^7^ CFU/mL, equally divided to 2 culture tubes of 15 mL (3 mL in each tube) and incubated for 46 h. Following incubation, supernatants from each two duplicate tubes were combined and harvested by pelleting bacterial cells at 3200 *g* at 4 °C for 30 min, followed by filtration through 0.2 μ cellulose acetate filters. Larger molecules in the supernatant were then concentrated by transferring to Amicon® Ultra-4 10 KDa filters and centrifugation at 3200 *g* at 4 °C for 60 min. Samples were then placed on ice until use for infection experiments in liquid culture and frozen at −20 °C prior to supernatant analysis by protein MS.

### Supernatant analysis by protein MS

Supernatant samples were subjected to in-solution tryptic digestion following the S-trap protocol (Protifi) followed by a solid phase extraction cleanup step. The resulting peptides were analyzed using nanoflow liquid chromatography (nanoAcquity) coupled to high resolution, high mass accuracy mass spectrometry (Thermo Q-Exactive HFX). The samples were analyzed on the instrument separately in a random order in discovery mode. Raw data was processed with the MetaMorpheus v1.0.2 informatics platform. The data were searched with semi-tryptic parameters against a protein database containing the *Streptomyces coelicolor* protein database as downloaded from Uniprot (Bateman et al., 2023) on August 2023, the WT and mutant *Salinispora* protease, and predicted phage Alon open reading frames larger than 100 amino acids extracted using NCBI ORFFinder (Sayers et al., 2022), and a list of common lab contaminants.

### Infection experiments in liquid cultures with exogenously added supernatant

Spores of *Streptomyces coelicolor* M1146 not encoding WT *Salinispora* protease (Table S3, strain 2) were added to 750 μL of DNB MMC in 9 culture tubes of 15 mL at a concentration of 3×10^7^ CFU/mL, along with phage Alon at MOI=0.1 and 10 μL of harvested supernatants from strains 5-7 (Table S3) in biological triplicates and incubated for 48 h. Following incubation phages were harvested by pelleting bacterial cells at 3200 *g* at 4 °C for 30 min, followed by filtration through 0.2 μ cellulose acetate filters. Phage titer was then measured by plaque assay.

### Selection for phages that escape the antiphage activity of the *Salinispora* protease

Three biological replicates of each phage (Alon or Tapuz) were amplified from single plaques on *S. coelicolor* M145. Each biological replicate of the phage was amplified on *S. coelicolor* Strains 5 and 6 (Table S3) expressing the WT *Salinispora* protease or the catalytically dead S219A mutant, each in three biological replicates, as well. The phage amplification was set up similarly to the infection experiment of *S. coelicolor* with phages in liquid culture, but a higher volume and a variable incubation time was used, as described herby.

Round 1: Spores of *S. coelicolor* encoding WT *Salinispora* protease and mutant S219A *Salinispora* protease (Table S3, strains 5-6) were added in biological triplicates to 1.5 mL of DNB MMC in 6 culture tubes of 15 mL at a concentration of 3×10^7^ CFU/mL, along with phage Alon or phage Tapuz at MOI=0.1 and incubated for 39 h. Following incubation, phages were harvested by pelleting bacterial cells at 3200 *g* at 4 °C for 30 min, followed by filtration through 0.2 μ cellulose acetate filters and phage titer was then measured by plaque assay.

Round 2: for phage Alon or phage Tapuz, three phage stocks that were amplified over Round 1 on *S. coelicolor* expressing WT *Salinispora* protease were added in 1/10 of the total volume into 1.5 mL of DNB MMC in 6 culture tubes (3 of *S. coelicolor* expressing WT *Salinispora* protease and 3 expressing mutant protease) at a concentration of 3×10^7^ CFU/mL. The amplification on *S. coelicolor* expressing WT *Salinispora* proteas was used for escaper isolation while the amplification over on *S. coelicolor* expressing mutant S219A *Salinispora* protease was used as a control to assess escaper phenotype evolution. MOI was varied depending on the phage stock and ranged between 0.001 and 0.1. As a control, one phage stock amplified over Round 1 on *S. coelicolor* expressing mutant *Salinispora* protease (S219A) was added in MOI 0.1 and 0.001 for phage Alon and Tapuz, respectively, to two tubes with spores of *S. coelicolor* expressing WT and S219A mutant *Salinispora* protease at a concentration of 3×10^7^ CFU/mL. All tubes were incubated for 50 h. Following incubation, phages were harvested by pelleting bacterial cells at 3200 *g* at 4 °C for 30 min, followed by filtration through 0.2 μ cellulose acetate filters and phage titer was then measured by plaque assay.

Round 3: for phage Alon or phage Tapuz, three phage stocks amplified over Round 2 on *S. coelicolor* expressing WT *Salinispora* protease were added in 0.1 and 0.01 MOI for phage Alon and Tapuz, respectively, into 1.5 mL of DNB MMC in 6 culture tubes (3 of *S. coelicolor* expressing WT *Salinispora* protease and 3 expressing mutant protease) at a concentration of 3×10^7^ CFU/mL. As a control, one phage stock amplified over Round 2 on *S. coelicolor* expressing mutant *Salinispora* protease (S219A) was added in MOI 0.1 to two tubes with spores of *S. coelicolor* expressing WT and S219A mutant *Salinispora* protease at a concentration of 3×10^7^ CFU/mL.

For each of Round 2 and Round 3, phages that were amplified in liquid culture on *S. coelicolor* expressing WT *Salinispora* protease were plated to obtain single plaques on *S. coelicolor* expressing WT *Salinispora* protease. Control phages that were amplified in liquid culture on *S. coelicolor* expressing S219A mutant protease were plated on *S. coelicolor* expressing S219A mutant *Salinispora* protease. Except for the bacterial strain expressing *Salinispora* protease, single plaque plating was as described above: phage stocks were serially diluted and added to a 100 μL spore suspension of *S. coelicolor* (expressing WT or S219A *Salinispora* protease at 10^8^ CFU / mL) in 15 mL screw cap tubes. 5 mL of DNB MMC with 0.5% agar at 50 °C were then added and the resulting phage/spore suspension was mixed and plated on 100 × 15 mm plates with 20 mL DNB 2% agar. The plates were incubated overnight at 30 °C. Single plaques were picked with a pipette tip into 120 μL phage buffer.

Half of the single plaque suspension was used for infection in liquid culture with *S. coelicolor* expressing WT *Salinispora* protease and half was used for infection in liquid culture with *S. coelicolor* expressing S219A mutant *Salinispora* protease at a spore concentration of 3×10^7^ CFU/mL. Total volume in DNB MMC was 1.5 mL and incubation was for two days. Following incubation, phages were harvested by pelleting bacterial cells at 3200 *g* at 4 °C for 30 min, followed by filtration through 0.2 μ cellulose acetate filters and phage titer was then measured by plaque assay. Phage DNA was harvested from single plaque thus amplified in liquid culture on *S. coelicolor* expressing WT protease if the titer was <l0x reduced compared with the same single plaque suspension amplified on *S. coelicolor* expressing mutant S219A *Salinispora* protease. Such escaper phenotype was markedly different than the >100X titer reduction commonly observed for WT phages Alon and Tapuz (Figure S1). Phage DNA sequencing and mutation analysis was performed as previously reported (Stokar-Avihail et al., 2023).

### Phage purification by CsCl gradient

Phages Alon, Dagobah and Kromp were amplified on 3 plates of 150 × 15 mm dimensions each of *S. coelicolor* M145 to obtain 40 mL stocks of 1×10^11^ PFU/mL (Alon and Dagobah) and 2×10^10^ PFU/mL (Kromp). Phage purification was as previously reported (Hör et al., 2024), with modifications as described hereby. Phages were pelleted at 25,000 *g* at 4 °C for 2 h. The pellet was resuspended with 0.5 mL of phage buffer and loaded on CsCl step gradients (2 ml of ρ = 1.3, 3 ml of ρ = 1.4, 3 ml of ρ = 1.5, 2 ml of ρ = 1.7; all in phage buffer) formed in open-top polyclear ultracentrifugation tubes (Seton Scientific). The gradients were centrifuged in an SW41 rotor (Beckman) at 25,000 rpm and 4 °C for 2 h and the phage bands collected by needle side-puncture. Buffer exchange for the extracted phages was done using Amicon 4 mL centrifugal filters (10 kDa molecular weight cutoff) in steps reducing ionic strength. First, 3 M NaCl in phage buffer was added followed by 3200 g at 4 °C for 15 min and the flowthrough was removed. 2 M NaCl in phage buffer was then added followed by 3200 g at 4 °C for 15 min and the flowthrough was removed. 1 M NaCl in phage buffer was then added followed by 3200 g at 4 °C for 15 min and the flowthrough was removed. Phage buffer was then added followed by 3200 *g* at 4 °C for 15 min and the flowthrough was removed. Finally, phage buffer was added followed by 3200 *g* at 4 °C for 60 min for a final purified phage volume of 100 μL.

### Purified phage incubation with concentrated supernatant

Purified phage, phage buffer and concentrated supernatant were mixed in PCR tubes as follows: 4 μL of purified phage (Alon, Dagobah or Kromp, purified as described in the section above discussing CsCl gradient), 2 μL of concentrated supernatant (from *S. coelicolor* expressing WT *Salinispora* protease or mutant S219A *Salinispora* protease, each in three biological replicates; harvested as described in the section titled ‘supernatant harvest from *S. coelicolor* cultures’) and 2 μL of phage buffer. Samples were incubated without shaking at the following temperature gradient: 3 h at 30 °C, 3 h at 25 °C, 3 h at 20 °C, 3 h at 15 °C, 7 h at 10 °C and 7 h at 4 °C. Phage titers were counted by plaque assay as described above. Complete phage particles with and without DNA were counted from electron microscopy images following negative staining as described below.

### Negative staining and electron microscopy imaging

For negative staining, formvar and carbon-coated copper grids (300 mesh, Electron Microscopy Sciences) were glow discharged in an Evactron CombiClean Decontaminator (XEI Scientific) for 2 min at 0.067 mbar (air) and 18 W. 3 μL of the phage sample was then directly applied to the grid, incubated for 1 min and blotted with filter paper (Whatman Grade 1). The grid was briefly washed on a drop of water and blotted with filter paper. Staining was performed by touching a 10 μL drop of 2% uranyl acetate, followed by blotting with filter paper. This step was repeated once more. Then the grid was placed on a third 10 μL drop of uranyl acetate (2%) for 1 min. Finally, the grid was blotted with filter paper and air dried. Imaging was performed using a Tecnai Spirit (Bio Twin) transmission electron microscope (Thermo Fisher Scientific) at an accelerating voltage of 120 kV and a Gatan OneView camera.

## Acknowledgements

We thank members of the Sorek lab for constructive discussion. We thank the Bar-Shir lab and the Stern-Ginossar lab for sharing their facilities for conducting this research. R.S. was supported, in part, by the European Research Council (grant ERC-AdG GA 101018520), the Israel Science Foundation (MAPATS grant 2720/22), the Deutsche Forschungsgemeinschaft (SPP 2330, grant 464312965), the Minerva Foundation with funding from the Federal German Ministry for Education and Research, a research grant from the Estate of Hermine Miller, the Institute for Environmental Sustainability (IES) and the Center for Immunotherapy at the Weizmann Institute of Science, and the Knell Family Center for Microbiology. E.H. was supported by the Israel Cancer Research Fund Postdoctoral Fellowship.

## Notes

### Competing Interest Statement

R.S. is a scientific cofounder and advisor of BiomX and Ecophage.

## References

1. Barbuto Ferraiuolo, S., Cammarota, M., Schiraldi, C., and Restaino, O.F. (2021). Streptomycetes as platform for biotechnological production processes of drugs. Applied Microbiology and Biotechnology 105, 551–568.

2. Bateman, A., Martin, M.-J., Orchard, S., Magrane, M., Ahmad, S., Alpi, E., Bowler-Barnett, E.H., Britto, R., Bye-A-Jee, H., Cukura, A., et al. (2023). UniProt: the Universal Protein Knowledgebase in 2023. Nucleic Acids Research 51, D523–D531.

3. Castillo, D., Rørbo, N., Jørgensen, J., Lange, J., Tan, D., Kalatzis, P.G., Svenningsen, S.L., and Middelboe, M. (2019). Phage defense mechanisms and their genomic and phenotypic implications in the fish pathogen Vibrio anguillarum. FEMS Microbiol Ecol 95.

4. Doron, S., Melamed, S., Ofir, G., Leavitt, A., Lopatina, A., Keren, M., Amitai, G., and Sorek, R. (2018). Systematic discovery of antiphage defense systems in the microbial pangenome. Science 359.

5. Eckhard, U., Schönauer, E., Nüss, D., and Brandstetter, H. (2011). Structure of collagenase G reveals a chew-and-digest mechanism of bacterial collagenolysis. Nat Struct Mol Biol 18, 1109–1114.

6. Figurski, D.H., and Helinski, D.R. (1979). Replication of an origin-containing derivative of plasmid RK2 dependent on a plasmid function provided in trans. Proceedings of the National Academy of Sciences 76, 1648–1652.

7. Gao, L., Altae-Tran, H., Böhning, F., Makarova, K.S., Segel, M., Schmid-Burgk, J.L., Koob, J., Wolf, Y.I., Koonin, E.V., and Zhang, F. (2020). Diverse enzymatic activities mediate antiviral immunity in prokaryotes. Science 369, 1077–1084.

8. Georjon, H., and Bernheim, A. (2023). The highly diverse antiphage defence systems of bacteria. Nature Reviews Microbiology 21, 686–700.

9. Gilchrist, C.L.M., Chooi, Y.-H., and Robinson, P. (2021). clinker & clustermap.js: automatic generation of gene cluster comparison figures. Bioinformatics 37, 2473–2475.

10. Gomez-Escribano, J.P., and Bibb, M.J. (2011). Engineering Streptomyces coelicolor for heterologous expression of secondary metabolite gene clusters. Microbial Biotechnology 4, 207–215.

11. Hardy, A., Sharma, V., Kever, L., and Frunzke, J. (2020). Genome Sequence and Characterization of Five Bacteriophages Infecting Streptomyces coelicolor and Streptomyces venezuelae: Alderaan, Coruscant, Dagobah, Endor1 and Endor2. Viruses 12, 1065–1079.

12. Hoque, M.M., Naser, I.B., Bari, S.M.N., Zhu, J., Mekalanos, J.J., and Faruque, S.M. (2016). Quorum Regulated Resistance of Vibrio cholerae against Environmental Bacteriophages. Scientific Reports 6.

13. Hör, J., Wolf, S.G., and Sorek, R. (2024). Bacteria conjugate ubiquitin-like proteins to interfere with phage assembly. Nature 631, 850–856.

14. Hurst, L.C., Badalamente, M.A., Hentz, V.R., Hotchkiss, R.N., Kaplan, F.T.D., Meals, R.A., Smith, T.M., and Rodzvilla, J. (2009). Injectable Collagenase Clostridium Histolyticum for Dupuytren’s Contracture. New England Journal of Medicine 361, 968–979.

15. Jumper, J., Evans, R., Pritzel, A., Green, T., Figurnov, M., Ronneberger, O., Tunyasuvunakool, K., Bates, R., Žídek, A., Potapenko, A., et al. (2021). Highly accurate protein structure prediction with AlphaFold. Nature 596, 583–589.

16. Kato, J.-y., Chi, W.-J., Ohnishi, Y., Hong, S.-K., and Horinouchi, S. (2005). Transcriptional Control by A-Factor of Two Trypsin Genes in Streptomyces griseus. Journal of Bacteriology 187, 286–295.

17. Kim, I.S., and Lee, K.J. (1995). Physiological roles of leupeptin and extracellular proteases in mycelium development of Streptomyces exfoliatus SMF13. Microbiology 141, 1017–1025.

18. Kronheim, S., Daniel-Ivad, M., Duan, Z., Hwang, S., Wong, A.I., Mantel, I., Nodwell, J.R., and Maxwell, K.L. (2018). A chemical defence against phage infection. Nature 564, 283–286.

19. Leforestier, A., and Livolant, F. (2010). The Bacteriophage Genome Undergoes a Succession of Intracapsid Phase Transitions upon DNA Ejection. Journal of Molecular Biology 396, 384–395.

20. Luthe, T., Kever, L., Hänsch, S., Hardy, A., Tschowri, N., Weidtkamp-Peters, S., and Frunzke, J. (2023). Streptomyces development is involved in the efficient containment of viral infections. microLife 4, 1–13.

21. Manteca, A., Alvarez, R., Salazar, N., Yagüe, P., and Sanchez, J. (2008). Mycelium Differentiation and Antibiotic Production in Submerged Cultures of Streptomyces coelicolor. Applied and Environmental Microbiology 74, 3877–3886.

22. Matthews, B.W., Sigler, P.B., Henderson, R., and Blow, D.M. (1967). Three-dimensional Structure of Tosyl-α-chymotrypsin. Nature 214, 652–656.

23. Millman, A., Melamed, S., Leavitt, A., Doron, S., Bernheim, A., Hör, J., Garb, J., Bechon, N., Brandis, A., Lopatina, A., et al. (2022). An expanded arsenal of immune systems that protect bacteria from phages. Cell Host & Microbe 30, 1556–1569.e1555.

24. Mordret, E., Hervé, A., Vaysset, H., Clabby, T., Tesson, F., Shomar, H., Lavenir, R., Cury, J., and Bernheim, A. (2025). Protein and genomic language models chart a vast landscape of antiphage defenses. bioRxiv.

25. Morrison, H. (2021). Chymotrypsin. In Enzyme Active Sites and their Reaction Mechanisms, pp. 41–44.

26. Ramírez-Larrota, J.S., and Eckhard, U. (2022). An Introduction to Bacterial Biofilms and Their Proteases, and Their Roles in Host Infection and Immune Evasion. Biomolecules 12, 306–324.

27. Román-Ponce, B., Millán-Aguiñaga, N., Guillen-Matus, D., Chase, A.B., Ginigini, J.G.M., Soapi, K., Feussner, K.D., Jensen, P.R., and Trujillo, M.E. (2020). Six novel species of the obligate marine actinobacterium Salinispora, Salinispora cortesiana sp. nov., Salinispora fenicalii sp. nov., Salinispora goodfellowii sp. nov., Salinispora mooreana sp. nov., Salinispora oceanensis sp. nov. and Salinispora vitiensis sp. nov., and emended description of the genus Salinispora. International Journal of Systematic and Evolutionary Microbiology 70, 4668–4682.

28. Russell, D.A., Hatfull, G.F., and Wren, J. (2017). PhagesDB: the actinobacteriophage database. Bioinformatics 33, 784–786.

29. Sayers, Eric W., Beck, J., Bolton Evan E., Brister, J R., Chan, J., Connor, R., Feldgarden, M., Fine Anna M., Funk, K., Hoffman, J., et al. (2025). Database resources of the National Center for Biotechnology Information in 2025. Nucleic Acids Research 53, D20–D29.

30. Sayers, E.W., Bolton, E.E., Brister, J.R., Canese, K., Chan, J., Comeau Donald C., Connor, R., Funk, K., Kelly, C., Kim, S., et al. (2022). Database resources of the national center for biotechnology information. Nucleic Acids Research 50, D20–D26.

31. Schrodinger, LLC (2015). The PyMOL Molecular Graphics System, Version 1.8.

32. Shepherd, M.D., Kharel, M.K., Bosserman, M.A., and Rohr, J. (2010). Laboratory Maintenance of Streptomyces Species. Curr Protoc Microbiol 18.

33. Shomar, H., Tesson, F., Guillaume, M., Ongenae, V., Le Bot, M., Georjon, H., Mordret, E., Zhang, L., van Wezel, G.P., Rozen, D., et al. (2024). A family of lanthipeptides with anti-phage function. bioRxiv.

34. Stokar-Avihail, A., Fedorenko, T., Hör, J., Garb, J., Leavitt, A., Millman, A., Shulman, G., Wojtania, N., Melamed, S., Amitai, G., et al. (2023). Discovery of phage determinants that confer sensitivity to bacterial immune systems. Cell 186, 1863–1876.e1816.

35. Tal, N., and Sorek, R. (2022). SnapShot: Bacterial immunity. Cell 185, 578–578.e571.

36. Teufel, F., Almagro Armenteros, J.J., Johansen, A.R., Gíslason, M.H., Pihl, S.I., Tsirigos, K.D., Winther, O., Brunak, S., von Heijne, G., and Nielsen, H. (2022). SignalP 6.0 predicts all five types of signal peptides using protein language models. Nature Biotechnology 40, 1023–1025.

37. van Kempen, M., Kim, S.S., Tumescheit, C., Mirdita, M., Lee, J., Gilchrist, C.L.M., Söding, J., and Steinegger, M. (2023). Fast and accurate protein structure search with Foldseek. Nature Biotechnology 42, 243–246.

38. Yousef, G.M., Elliott, M.B., Kopolovic, A.D., Serry, E., and Diamandis, E.P. (2004). Sequence and evolutionary analysis of the human trypsin subfamily of serine peptidases. Biochim Biophys Acta 1698, 77–86.

39. Zhang, J.J., Tang, X., and Moore, B.S. (2019a). Genetic platforms for heterologous expression of microbial natural products. Natural Product Reports 36, 1313–1332.

40. Zhang, J.J., Yamanaka, K., Tang, X., and Moore, B.S. (2019b). Direct cloning and heterologous expression of natural product biosynthetic gene clusters by transformation-associated recombination. In Chemical and Synthetic Biology Approaches To Understand Cellular Functions - Part A, pp. 87–110.

41. Zheng, W., Wang, F., Taylor, N.M.I., Guerrero-Ferreira, R.C., Leiman, P.G., and Egelman, E.H. (2017). Refined Cryo-EM Structure of the T4 Tail Tube: Exploring the Lowest Dose Limit. Structure 25, 1436–1441.e1432.

